# Stoichiometric binding of Cyclophilin-A to the HIV-1 capsid modulates its mechanoelastic properties

**DOI:** 10.64898/2026.02.20.707123

**Authors:** Juan S. Rey, Alexander J. Bryer, Juan R. Perilla

**Affiliations:** Department of Chemistry & Biochemistry, University of Delaware, Newark, DE 19716

## Abstract

HIV-1 nuclear entry requires a capsid that is sufficiently stable to protect and transport the viral genome yet sufficiently deformable to traverse the nuclear pore complex. Cyclophilin A (CypA), a host cell factor that binds the capsid in the cytoplasm, regulates early replication events; however, its effects on capsid mechanics remain unresolved. Here, we used atomic force microscopy nanoindentation simulations of CypA-decorated HIV-1 capsids across binding stoichiometries to characterize their mechanical response. The capsid exhibits curvature-dependent mechanical heterogeneity, with stiffness and transverse deformability varying along its surface. CypA binding progressively increased capsid brittleness, promoting structural failure at lower deformations. At high CypA:CA ratios, CypA binding overrides intrinsic sequence-dependent differences in ductility across wild-type and mutant capsids. These findings establish a direct link between CypA binding and capsid mechanoelastic properties, supporting a stoichiometry-dependent model in which balanced CypA binding preserves flexibility for nuclear entry, whereas excessive binding compromises nuclear import.

## Introduction

The HIV-1 capsid, a protein shell composed of ∼200-250 capsid protein (CA) hexamers and exactly 12 CA pentamers[1, 2], is an essential factor at multiple stages of the viral replication cycle[3, 4, 5]. Following viral envelope fusion with the cell membrane, the viral core, consisting of the capsid enclosing the viral RNA genome and viral enzymes, is released into the host cell cytoplasm. The capsid works as a molecular vessel that engages host cell motor proteins to transport the viral genome through the cytosol toward the cellular nucleus[6, 7, 8, 9]. During this trafficking process, the capsid also serves as a reaction compartment for reverse transcription[10, 11], in which viral double-stranded DNA is copied from viral single-stranded RNA by reverse transcriptase (RT)[12]. Upon reaching the nuclear envelope, the core interacts with nucleoporins at the nuclear pore complex (NPC) and is imported into the nucleus[13, 14], where it uncoats to release the viral DNA and associated enzymes[15, 16, 17, 18], enabling integration of the viral genome into host cell chromatin[19]. Owing to its involvement in multiple steps of the retroviral replication cycle, the HIV-1 capsid constitutes an attractive target for antiretroviral drug development[20, 21, 22, 23].

HIV-1 replication critically depends on the direct interaction between the capsid and human host cell factors[24]. During cellular trafficking of the viral core, the capsid is bound by multiple host proteins, including Cyclophilin A (CypA) in the cytoplasm[25, 26], nucleoporins RANBP2 and NUP153 at the NPC[27, 28, 29], and cleavage and polyadenylation specificity factor 6 (CPSF6)[30] in the nucleus. Host factor CypA is a peptidyl-prolyl isomerase involved in T cell signal transduction, immunosuppression, and protein folding [31, 32, 33] that binds the HIV-1 CA via a proline-rich loop spanning residues 85 to 93 in the CA N-terminal domain (Figure 1)[34, 35]. CypA binding stabilizes and shields the capsid from antiretroviral restriction factors[36], and promotes reverse transcription, nuclear entry and integration in a cell line dependent manner[37, 27, 38, 25, 26, 39]. Moreover, the CypA-CA interaction modulates the rate of nuclear import[40], and several nucleoporin interactions with the capsid depend on CypA[41, 42] or engage the same CA binding site[27, 43].

**Figure 1:**
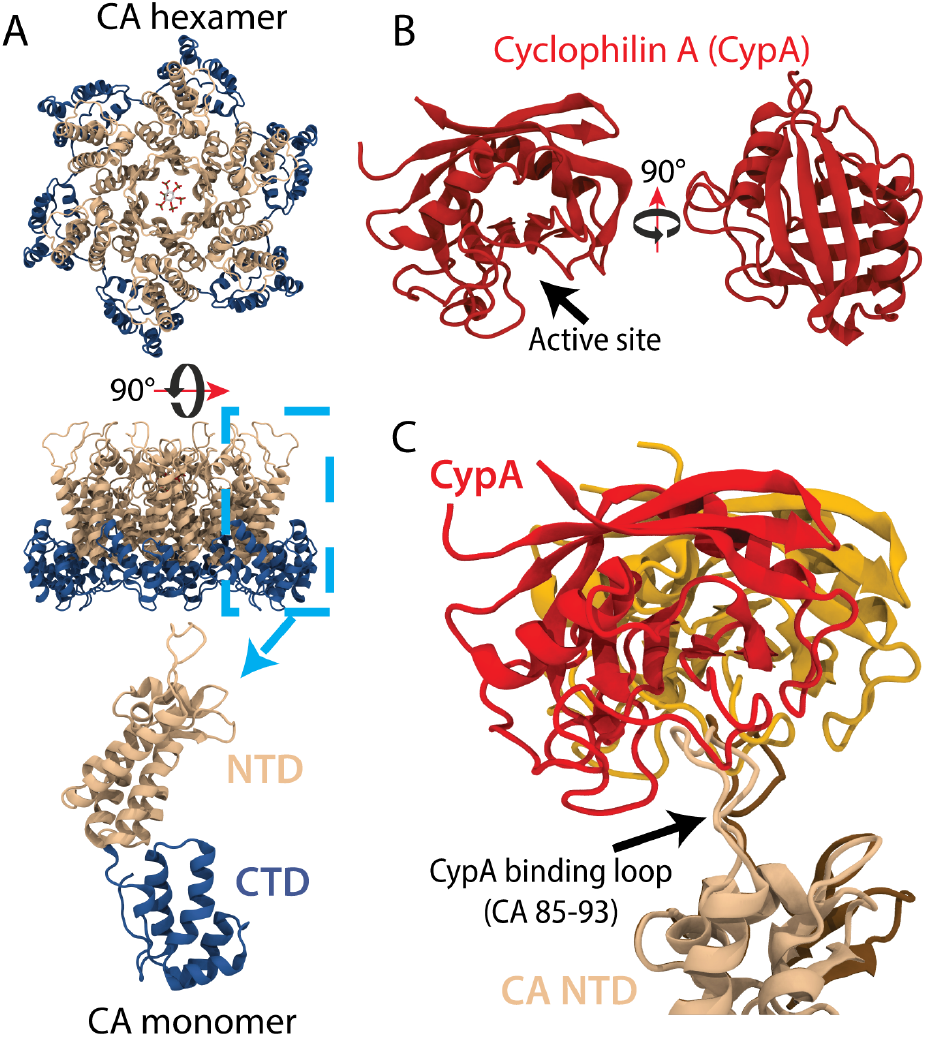
Cyclophilin A (CypA) is a human host factor that directly binds the HIV-1 capsid. (A) Top and side views of the HIV-1 capsid protein (CA) hexamer structure bound by assembly factor inositol hexakisphosphate (IP6) in cartoon representation. Dashed cyan inset depicts a CA monomer in cartoon representation, colored by domain: CA N-terminal domain (NTD) in tan; CA C-terminal domain (CTD) in dark blue; IP6 is shown in licorice representation and colored by atomic element (carbon in gray, oxygen in red, phosphate in yellow). (B) Cartoon representation of CypA crystal structure (PDBID 1AK4); front and side views. CypA active site is indicated by a black arrow. (C) Canonical CypA-CA binding interface engages a proline rich CA NTD loop spanning residues 85-93, termed the CypA-binding loop (black arrow). The two conformations of CypA and CA NTD from the crystal structure (PDBID 1AK4) are shown in cartoon representation and colored red/orange (CypA) and tan/brown (CA NTD).

In addition to host factor interactions, the intrinsic physical properties of the capsid play a critical role in nuclear entry. Using experimental and *in silico* atomic force microscopy (AFM) nanoindentation, we previously demonstrated that HIV-1 capsid elasticity is strongly correlated with nuclear entry and viral infectivity[44]. Hyperstable E45A CA mutant capsids[45], which are defective in nuclear entry and infectivity, were found to be significantly more brittle than wild-type (WT) capsids. In contrast, capsids carrying the compensatory R132T mutation[46] recovered WT-like elasticity and displayed near-WT levels of nuclear entry and infectivity. Consistent with these findings, recent cryo-electron microscopy studies have shown that the NPC selectively permits nuclear entry of small and elastic viral cores, while brittle cores fail to enter the nucleus[14]. These observations suggest a close relationship between the capsid’s mechanoelastic properties, i.e., its ability to undergo elastic deformation under applied mechanical force, and its biological function during HIV-1 replication.

Despite previous efforts, the exact mechanism by which CypA binding modulates nuclear entry remains unclear, and the effect of host factor binding on capsid mechanical properties is largely unexplored. Here, we extend our simulated AFM nanoindentation methodology to test the hypothesis that CypA binding modulates the mechanoelastic properties of the capsid, and thereby influences its ability to undergo nuclear entry. To this end, we developed a methodology to construct models of CypA-decorated capsids across biologically relevant CypA:CA stoichiometries. Our simulations reveal that the capsid behaves as a mechanically heterogeneous material with distinct deformation regimes. In the elastic regime, capsid stiffness was not correlated with the amount of bound CypA. In contrast, under fast nanoindentations probing the ductile regime and mechanical failure, we observe strong correlations between compressive strength, critical strain, and the CypA:CA bound fraction, indicating that capsid brittleness is increased in proportion with the amount of CypA bound. Although E45A/R132T revertant capsids recovered near-WT ductility relative to the the brittle E45A capsids at low amounts of bound CypA, all capsids become excessively brittle at high CypA:CA stoichiometries. Overall, our findings support a model in which CypA binding fine-tunes capsid elasticity, thereby regulating nuclear entry.

## Methods

### Curvature-based modeling of CypA-decorated capsids

Models of HIV-1 capsids bound by CypA at varying CypA:CA stoichiometries, hereafter called CypA-decorated capsids, were generated using a curvature-aware random-walk algorithm informed by experimental structural data (Figure 2). Cryo-electron microscopy (Cryo-EM) structures of CypA bound to CA tubes[47, 48] and CypA-DsRed bound to conical viral cores[49] indicate that CypA preferentially binds along curved regions of the capsid surface and engages multiple CA monomers across adjacent capsomers. Capsid curvature was therefore quantified on an all-atom model of the HIV-1 capsid[49] by calculating the bite angle between adjacent capsomer pairs (Supplementary Figure 1A-B), and this metric was used to guide CypA placement.

**Figure 2:**
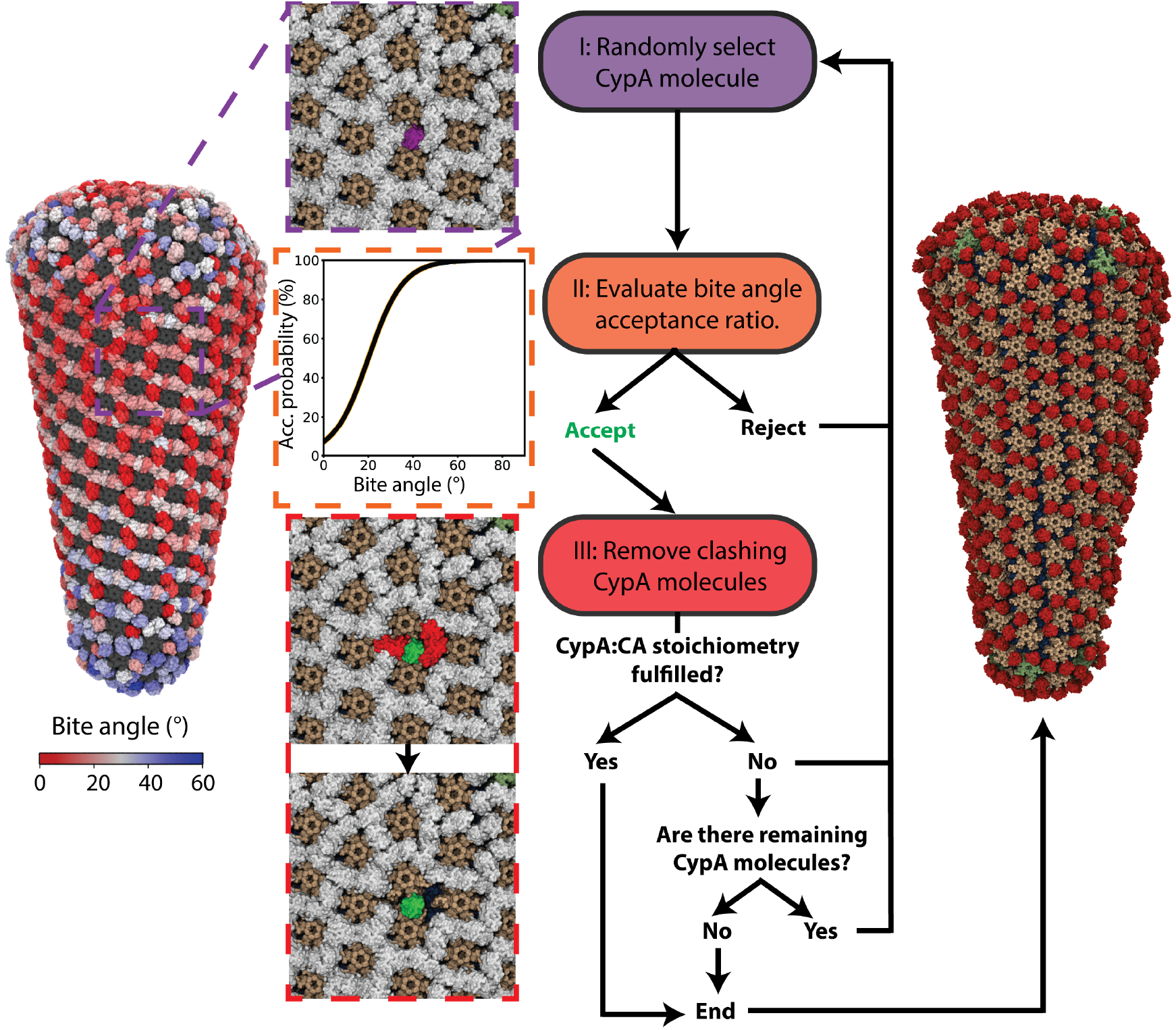
CypA-decorated capsid model construction using a curvature-aware random-walk algorithm. (Left) Fully CypA occupied capsid model (1:1 CypA:CA stoichiometry). Molecular surface of the capsid is shown in gray, and individual CypA molecules are colored according to the bite angle of the CA capsomers involved in the respective CypA binding site. (Center) Flowchart schematic illustrating the iterative selection or rejection of CypA molecules using a random-walk algorithm. At each iteration, (I) a CypA molecule is randomly selected from the fully occupied model and (II) accepted or rejected according to a curvature-weighted acceptance probability. (III) If accepted, sterically incompatible neighboring CypA molecules are removed. This process is repeated until the desired CypA:CA stoichiometry is reached or until all sterically consistent CypA molecules have been accepted. Steps I-III of the iterative procedure are indicated in purple, orange and red boxes, respectively. Molecular visualizations of these steps depict capsomers in tan (CA NTD) and blue (CA CTD), while CypA molecules are colored according to their state in the random-walk algorithm: unselected (white), randomly selected (purple), accepted (green) and removed (red). (Right) Resulting CypA-decorated capsid model after removal of all sterically incompatible CypA molecules, corresponding to an approximately 1:2.8: CypA:CA stoichiometry (fully saturated binding). The molecular surface is colored to distinguish CA NTD (tan), CA CTD (blue), CA pentamer NTD (green) and CypA (red).

An oversaturated CypA-decorated capsid model corresponding to a 1:1 CypA:CA stoichiometry was first constructed using an all-atom apo capsid structure[49] and the X-ray structure of the CypA-CA NTD complex (PDB ID: 1AK4)[35] (Figure 2, left). One of the two CypA-CA NTD binding poses (Figure 1D) was randomly selected and aligned with the CA NTD domain of each CA monomer in the capsid. Coordinates of the aligned CypA molecule and the corresponding CypA-binding loop were then merged with the apo capsid coordinates to generate an oversaturated capsid structure, in which each CA monomer is bound by CypA. Adjacent CypA molecules in the oversaturated CypA-decorated capsid model overlapped significantly, indicating that this stoichiometry is not structurally feasible[50, 47].

Sterically consistent CypA-decorated capsid models at lower stoichiometries were generated using a curvature-aware random-walk algorithm (Figure 2, center). Individual CypA molecules in the oversaturated CypA-decorated capsid model were randomly selected and accepted or rejected according to a Metropolis criterion[51] (Figure 2,center) based on local curvature, as quantified by bite angles. The acceptance probability P was defined as a sigmoid function

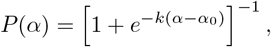

where α is the bite-angle of the CA capsomer pair forming the CypA binding site, α_0_ = 20^◦^ corresponds to the mean bite-angle observed in CA-CypA assemblies in CA tubes[47, 48], and 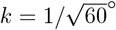 reflects the standard deviation of bite-angles across the capsid, in degrees. Upon acceptance of a CypA molecule, sterically overlapping neighboring CypA molecules were removed. This process was performed iteratively until the number of CypA molecules accepted matched a desired stoichiometry, or until no sterically consistent CypA molecule placements were possible.

The range of modeled CypA:CA stoichiometries was selected based on experimental observations. Fully saturated, sterically consistent CypA-decorated capsid models generated with the presented algorithm correspond to an approximate CypA:CA stoichiometry of 1:2.8, consistent with the experimentally observed maximum CypA:CA binding ratio (0.3-0.4) for both CA tubes [47] and conical capsids [50]. In addition, HIV-1 virions package CypA at an approximate stoichiometry of 1:10 CypA:CA[52, 53]. Based on these observations, we constructed CypA-decorated capsid models spanning a biologically relevant CypA:CA stoichiometry range between 1:3 to 1:10, as summarized in Supplementary Table 1. For each CypA:CA stoichiometry, an ensemble of ten independent CypA-decorated capsid models was generated, and the model with the highest cumulative acceptance probability was selected for subsequent simulations. The resulting CypA-decorated capsid models exhibited preferential CypA binding along curved regions of the capsid surface (Figure 2, right; Supplementary Figure 1D), with individual CypA molecules engaging up to three CA monomers from adjacent capsomers, consistent with previous high-resolution cryo-electron microscopy densities of CypA binding to CA tubes[48] and capsids[49].

### Shaped-based coarse-grained modeling of CypA

A shaped-based coarse-grained (SBCG2) model of CypA was derived following our previously developed methodology[54] (Figure 3). Briefly, CypA coarse-grained topologies were generated at varying levels of granularity, ranging from 5 to 500 beads in increments of 5 beads. The accuracy of each SBCG2 topology was evaluated by comparing its effective charge distribution to that of the all-atom CypA structure using Fourier Shell Correlation (FSC) [55] analysis (Figure 3A-B; Supplementary Figure 2). The SBCG2 topology that consistently achieved sub-nanometer effective charge resolution was selected for parameterization (Figure 3C). Based on the FSC criterion, we selected a SBCG2 CypA model comprising 165 beads, which yielded an effective charge resolution of 8.4 Å at a FSC threshold of 0.5 (Figure 3D).

**Figure 3:**
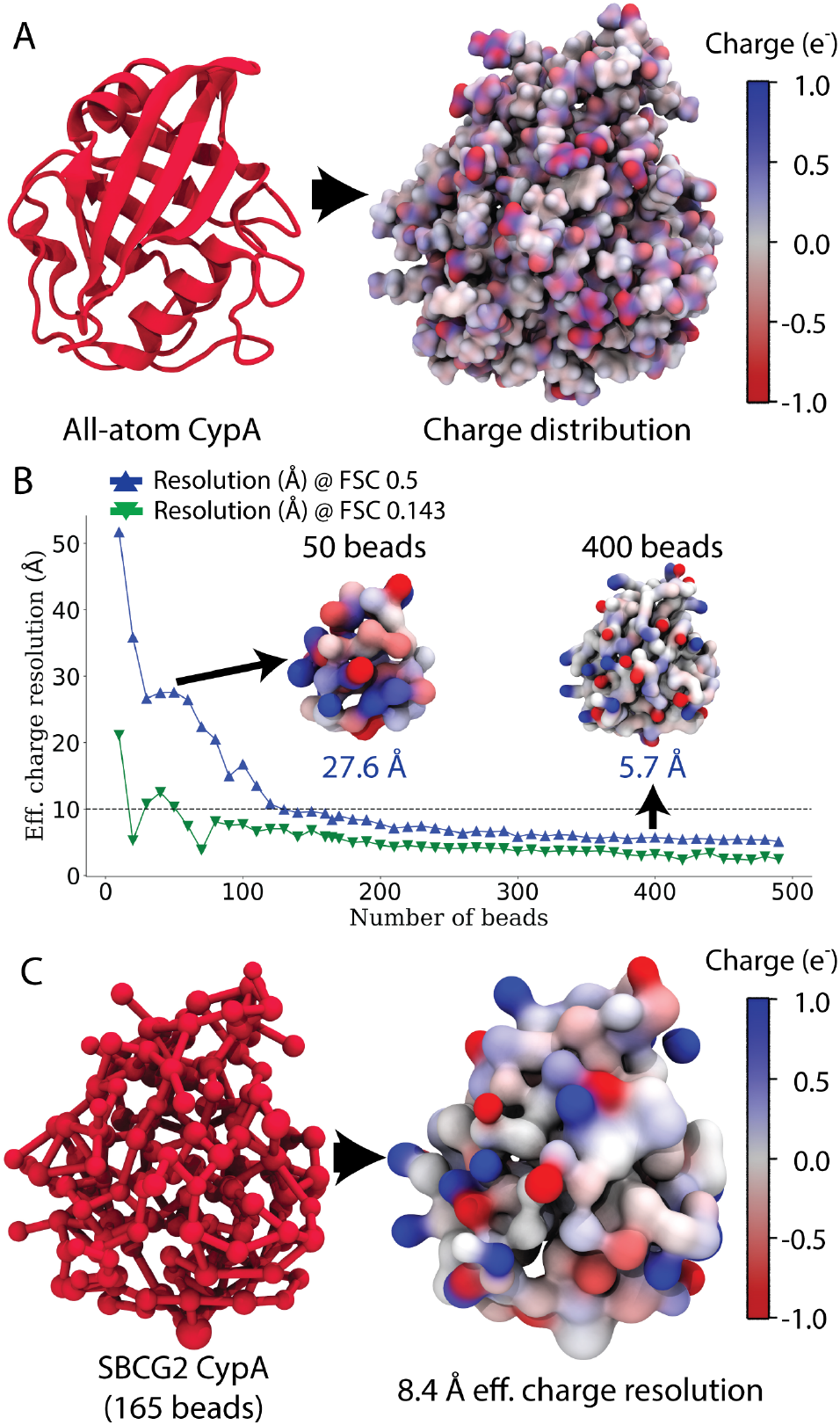
Shape-based coarse-grained (SBCG2) modeling of CypA. (A) Cartoon representation of the CypA all-atom (AA) structure (red) and its corresponding charge distribution map, colored by charge, the color scale uses the charge of the electron (e^−^) as a unit. (B) Effective charge resolution of CypA SBCG2 models generated across a range of granularities from 0 to 500 beads. Resolution is evaluated using Fourier shell correlation (FSC) between SBCG2 and AA charge distributions at FSC thresholds 0.5 (blue triangles) and FSC 0.143 (green inverted triangles). 1 nm effective charge resolution threshold is indicated with a dotted line. Example charge distribution maps for SBCG2 models with 50 and 400 beads are shown for illustration. (C) Ball-and-stick representation of the 165 bead CypA SBCG2 model used in this study (red) and its corresponding charge distribution map with 8.4 Å effective charge resolution. This model represents the lowest-granularity SBCG2 model that consistently captures the AA charge distribution at sub-nanometer resolution.

Bead connectivity in the SBCG2 models was derived from the atomic connectivity of the all-atom CypA structure. Initial bonds and angle parameters for the SBCG2 CypA model were obtained via Boltzmann inversion of all-atom molecular dynamics trajectories of a CypA-decorated CA trimer-of-dimers, following the legacy SBCG protocol [56]. These parameters were subsequently refined through an iterative procedure, in which, 20 ns simulations of the SBCG2 CypA model were performed, updated parameters were extracted via Boltzmann inversion of the SBCG2 trajectories, and the resulting parameters were compared against the all-atom reference, as described in the SBCG2 methodology [54]. An all-atom simulation of a CA trimer of dimers in complex with CypA was used as the reference during parameterization to implicitly include CypA dynamics in the context of the CA lattice into the SBCG2 model. In contrast to our previous SBCG2 parameterization of HIV-1 CA [54], pruning of angle parameters was not required, as the 165-bead SBCG2 CypA topology did not result in a hyperconnected network. The resulting parameterized SBCG2 CypA model accurately captures the charge distribution and conformational dynamics of the all-atom CypA structure, providing a robust coarse-grained representation suitable for simulations of CypA-decorated capsids.

The parameterized SBCG2 CypA model was utilized in conjunction with the previously derived SBCG2 model of HIV-1 CA[54] to map the all-atom CypA-decorated capsid models into a coarse-grained representation. The resulting SBCG2 CypA-decorated capsid models were then prepared for simulation by adding NaCl ions to a physiological concentration of 150mM using the *cionize* and *autoionize* plugins in Visual Molecular Dynamics (VMD)[57]. Explicit water solvent was not utilized, as solvent effects are implicitly incorporated into the SBCG2 parameterization procedure, which captures protein dynamics in aqueous environments [54]. The simulation ready SBCG2 CypA-decorated capsid systems contained between 366,880 to 424,906 beads depending on the number of CypA molecules bound, and encompassed SBCG2 models for the HIV-1 capsid, CypA molecules, the viral assembly cofactor IP6[58, 59] and ions. Compared to the corresponding all-atom systems, which comprise approximately 70 million atoms when solvated and ionized, the SBCG2 CypA-decorated capsid models substantially reduced system complexity and enabled efficient simulation of AFM nanoindentation.

### AFM nanoindentation simulations

In preparation for AFM nanoindentation simulations, CypA-decorated capsid coordinates were first optimized via energy minimization using a gradient descent algorithm for 25,000 steps, until the gradient converged below 1 kcal/mol/^Å2^. Each system was then heated under constant number of particles, volume and temperature (NVT) conditions, with the temperature increased gradually from 60 K to 298 K at a rate of 25 K/ns. The CypA-decorated capsids were subsequently equilibrated for over 2 µs in the NVT ensemble at 298 K, a timescale sufficient for convergence of the capsid volume and axial height (Supplementary Figure 3).

Following equilibration of the CypA-decorated capsids in isolation, each system was prepared in the AFM nanoindentation setup. Each CypA-decorated capsid was positioned above a flat plate composed of uniform SBCG beads with van der Waals (vdW) radii of 1 nm, and a spherical AFM tip of 13 nm diameter was placed above the capsid at one of four equidistant positions along its principal axis of inertia, as described previously[44]. These positions correspond to the broad end, upper half, lower half, and the narrow end of the capsid, referred to as tip positions 1 to 4 throughout the text (Figure 4A-D; Supplementary Figure 4A). The combined capsid-plate-tip system was then energy minimized, heated, and equilibrated following the protocol described above, to allow adsorption of the CypA-decorated capsid onto the AFM plate. During these preparation steps, the AFM plate and tip were restrained using harmonic positional restraints with a force constant of 10 kcal/mol/^Å2^.

**Figure 4:**
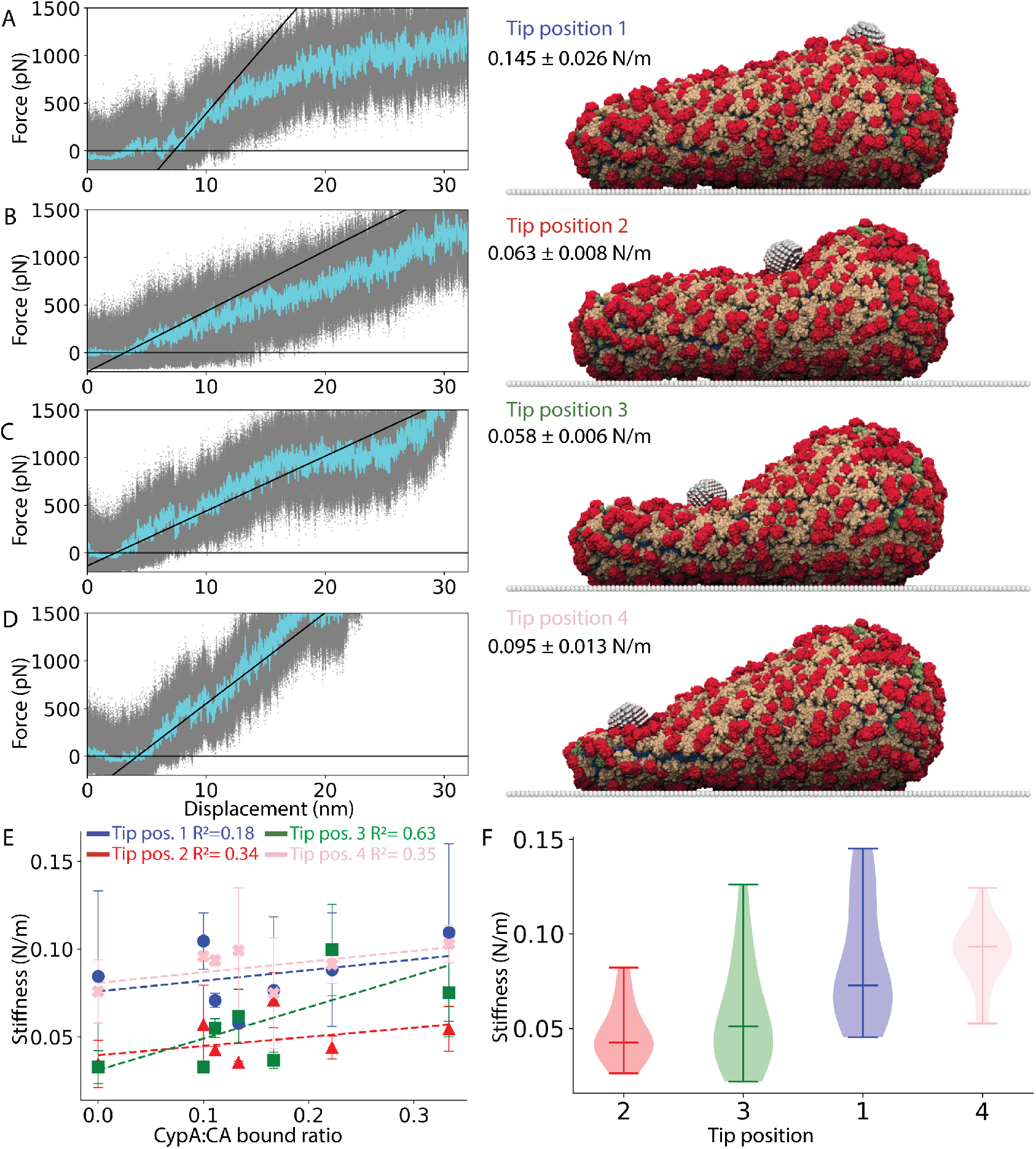
CypA-decorated capsid stiffness measured by *in silico* AFM nanoindentation. (A-D) Representative forcedisplacement profiles from AFM nanoindentation of a CypA-decorated capsid at 1:3 CypA:CA stoichiometry at four locations along the capsid: (A) the broad end (tip position 1), the flat regions in (B) the upper half (tip position 2) and (C) lower half (tip position 3), and (D) the narrow end (tip position 4). Raw force-displacement data points are shown in gray, with windowed-average traces are shown in cyan. Black solid lines represent linear fits to the first 3 nm of nanoindentation used to calculate capsid stiffness. Representative simulation snapshots of the capsids during nanoindentation are shown for each tip position, with the corresponding stiffness values reported in N/m. (E) Scatter plot for mean capsid stiffness as a function of the CypA:CA bound ratio across all tip positions. Dotted lines represent linear fitting of the data, with respective *R*^2^ values indicated. Error bars represent standard deviation from three independent simulation replicas (n=3). (F) Violin plots of stiffness distributions measured at each tip position, ordered by increasing local curvature. Distributions encompass all stiffness measurements at a tip position independent of CypA:CA stoichiometry (n=21). Solid lines indicate median, minimum and maximum values.

Nanoindentation simulations were performed by driving the AFM tip toward the capsid at a constant velocity using steered molecular dynamics (SMD) [60, 61, 62] as implemented in NAMD 3[63]. Nanoindentation simulations with an AFM tip velocity of 3.125 nm/µs, called slow nanoindentation simulations throughout the text (Supplementary Table 2, ID 1-7), were used to probe the elastic loading response of the capsid. In addition, fast nanoindentation simulations, with an AFM tip velocity of 312.5 nm/µs (Supplementary Table 2, ID 8-28), were performed to probe the ductile response of the capsid and induce mechanical failure. For each CypA:CA stoichiometry and tip position, three independent replicas of slow nanoindentation were conducted for 20 µs each. For each CypA:CA stoichiometry, tip position and CA sequence variant (WT, E45A, E45A/R132T), five independent replicas of fast nanoindentation were performed for 250 ns, which was a sufficient timescale to capture capsid structural failure events, as summarized in Supplementary Table 2.

In all simulations, temperature was controlled using a Langevin thermostat with a damping constant of 2 ps^−1^, with the AFM tip excluded from thermostat coupling to prevent stochastic thermal forces from perturbing the imposed constant velocity motion. Simulations were performed using a timestep of 40 fs for WT and E45A CA systems, and 30 fs for E45A/R132T CA systems. Short-range nonbonded interactions were calculated using a 20 Å cutoff and a switching distance of 18 Å, while long-range electrostatic interactions were calculated every step using the particle-mesh Ewald (PME) algorithm [64] with a grid spacing of 2 Å. Force and position vectors of the AFM tip during SMD simulations are reported every 100 steps. All simulations were performed using the NAMD molecular dynamics engine (version 3 alpha 13)[63], employing the GPU-resident mode for heating, equilibration and AFM nanoindentation simulations, and CPU-only mode for energy minimization [65].

### Post-processing of force-profile curves

Force-displacement profiles were extracted from each AFM nanoindentation simulation. The applied force F was calculated as the magnitude of the vector force exerted by the AFM tip, and the indentation displacement d was computed as the projection of the AFM tip displacement along the nanoindnetation axis 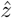 relative to its initial position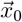:

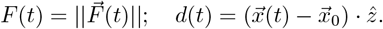

Force-displacement traces were smoothed using a windowed average of 1000 points and plotted together with the raw data points (Figure 4A-D).

Stiffness was quantified as the resistance to elastic deformation under small applied loads and is treated as an effective measure of the elastic (Young’s) modulus. Capsid stiffness was calculated from the force-displacement profiles extracted from slow AFM nanoindentation simulations by performing a linear fit over the first 3 nm of indentation, as described previously[44]. For each simulation condition, stiffness values were computed across three simulation replicates and reported as mean *±* standard deviation in units of N/m. Relationships between capsid stiffness, CypA:CA stoichiometry, and local curvature were evaluated using simple linear regression and Spearman rank-order correlation.

The Poisson’s ratio of the capsid was quantified as the negative ratio of transverse strain to axial strain during nanoindentation, reflecting the extent of lateral expansion accompanying axial compression[66]. This constitutes and independent mechanical metric that, together with stiffness characterizes the threedimensional elastic response of the capsid. To compute Poisson’s ratio, the local capsid height (axial dimension) and width (transverse dimension) within the compressed region were measured from the simulation trajectories. The percentage change in height and width were defined as the axial and transverse strains, respectively. Poisson’s ratio was calculated by linear fitting of the transverse versus axial strain curves, yielding the ratio of transverse expansion strain to axial compressive strain.

For fast nanoindentation simulations, force-displacement profiles were converted to stress-strain profiles by normalizing the applied force by the AFM tip-capsid contact surface area and the indentation displacement by the initial capsid height. Stress-strain traces were then smoothed using a windowed average of 1000 points to facilitate identification of mechanical regimes. The yield point, defined as the transition point from linear elastic to plastic deformation, was identified using the 0.2% offset yield criterion[67], while the critical point, right before failure occurs, was identified as the point of maximum stress preceding a sustained decrease greater than background stress fluctuations. A loading regime is defined as the region from indentation onset to the yield point, a ductile deformation regime as the region between the yield and critical points, and the failure regime as the region following the critical point. Compressive strength and critical strain were defined as the stress and strain at the critical point, that is, the maximum stress sustained prior to structural failure and the corresponding strain. Analogously, yield strain and stress were defined as the stress and strain values measured at the yield point. Ductility, defined as the ability of a material to undergo substantial deformation before failure, was quantified by the critical strain. Furthermore, Brittleness, which denotes mechanical failure upon compression after very little or no deformation, was characterized by increased compressive strength coupled with reduced critical strain values[66].

Mechanical regimes were further validated by visual inspection of the AFM nanoindentation trajectories and by quantifying local capsid deformation. Changes in capsid morphology were assessed by calculating the variations in the bite angle between adjacent capsomer pairs relative to the capsid before nanoindentation. Force-time and bite-angle traces for capsomers in contact with the AFM tip were used to identify the initial loading regime, characterized by increasing force with negligible curvature change, followed by a ductile regime, in which applied force induces increase in the local curvature, and a failure regime, where force decreases without further changes in average capsid curvature (Supplementary Figure 6). Yield strain, yield stress, critical strain, and compressive strength were calculated from each stress-strain profile, and mean and standard deviation values were computed across five independent simulations for each CypA:CA stoichiometry, tip position, and CA sequence condition. Correlations between mechanical parameters with the CypA:CA bind ratio were evaluated using simple linear regression, and goodness of fit was quantified by the coefficient of determination R^2^.

## Results

### Cyclophilin A binding does not substantially alter the stiffness of the HIV-1 capsid

We previously characterized the stiffness of the HIV-1 capsid using experimental and simulated AFM nanoindentation of coarse-grained HIV-1 capsid models[44]. To characterize the effect of CypA binding on capsid stiffness, we built SBCG2 models of CypA-decorated capsids and quantified their elastic loading response under small deformations using slow AFM nanoindentation simulations (3.125 nm/µs tip velocity; Supplementary Table 2, ID 1-7). Force-displacement curves were obtained for CypA-decorated capsid nanoindentation at four distinct positions along the capsid’s principal axis of inertia, starting from the broad end to the narrow end (Figure 4A-D). For each indentation position and CypA:CA stoichiometry, stiffness was calculated from the linear region of the force-displacement curve (see Methods). Across all indentation positions, capsid stiffness exhibited a weak positive correlation, with R^2^ 0.19-0.63, with the CypA:CA binding ratio (Figure 4E; Supplementary Figure 4B). Although mean stiffness values increased modestly with increased CypA:CA bound ratios, these weak correlations indicate that capsid stiffness is not strongly dependent on the amount of bound CypA. Consistent with previous observations that capsid stiffness is not correlated with nuclear entry or infectivity[44], these results suggest that CypA-mediated regulation of nuclear import is unlikely to be modulated by changes in capsid stiffness.

### The HIV-1 capsid is mechanically heterogeneous

We next tested the effect of the nanoindentation position on the measured capsid stiffness (Figure 4F). Indentation positions were ranked by the local curvature of the capsid lattice (i.e., 2<3<1<4): tip positions 2 and 3 lie along flatter regions of the capsid corresponding with a hexameric CA lattice, while tip positions 1 and 4 are located at the capsid ends which contain CA pentamers and have exhibit higher local curvature. An Alexander-Govern equality of the means test, revealed statistically significant differences among the mean stiffness at the four tip positions with p-value=1 × 10^−7^. Post-hoc pairwise comparisons between stiffness distributions using Tukey’s HSD test showed that mean stiffness values measured at the highly curved capsid ends (tip positions 1 and 4) were significantly different from those measured at the flatter regions of the capsid (tip positions 2 and 3). Spearman rank-order correlation analysis further demonstrated a monotonic increase in stiffness with local curvature of the nanoindentation position (ρ=0.62, p-value=5 × 10^−8^). These results are consistent with previous observations for capsids in the absence of CypA, where regions of higher curvature exhibited increased stiffness[44]. Thus, CypA-decorated capsids maintain the intrinsic curvature-stiffness relationship of the HIV-1 capsid.

During fast nanoindentation simulations (v = 312.5 nm/µs tip velocity), axial compression was accompanied by transverse expansion of the capsid. We quantified this response by estimating an effective Poisson’s ratio (ν) from the ratio of transverse to axial strain at tip positions 1 to 3 (Supplementary Figure 5). The Poisson’s ratio varied markedly with indentation location: compression at the highly curved broad end of the capsid resulted in minimal transverse expansion (ν=0.08), whereas indentations along flatter regions induced pronounced transverse expansions (ν ≈0.6). Although ν ≤0.5 for linear elastic materials, the higher Poisson’s ratio values observed at flatter regions reflect nonlinear inelastic response and volume changes in a coarsegrained nonequilibrium system under localized compressions at fast AFM tip velocities. Together, these findings further characterize the HIV-1 capsid as a mechanically heterogeneous structure whose mechanical response depends on the local CA lattice curvature.

**Figure 5:**
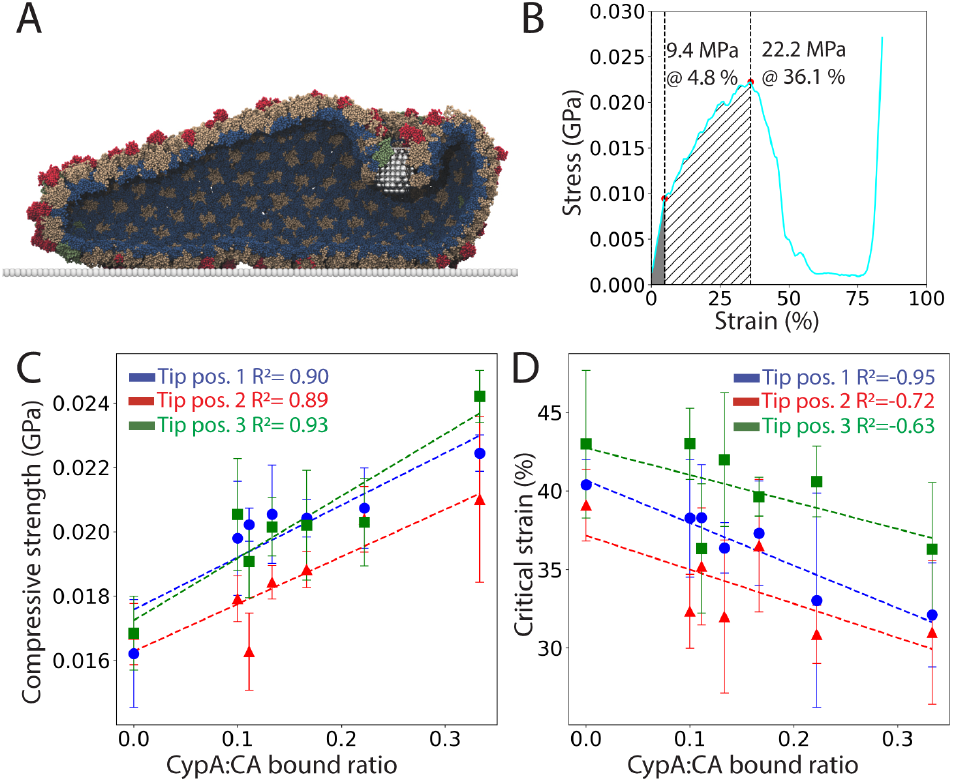
Characterization of capsid brittleness in fast *in silico* AFM nanoindentation simulations. (A) Representative simulation snapshot of a capsid failure event during fast AFM nanoindentation of a CypA-decorated capsid at 1:6 CypA:CA stoichiometry. During failure, the AFM tip disrupts the CA lattice and pierces through the capsid surface. SBCG2 model beads are shown using a van der Waals sphere representation, with CypA in red, CA hexamer NTD in tan, CA pentamer NTD in green, CA CTD in dark blue and the AFM tip and plate in metallic gray. One half of the capsid is removed to visualize the internal structure. (B) Representative stress-strain profile from fast AFM nanoindentation. An initial loading regime is observed (solid gray area) up to a yield point, followed by a ductile deformation regime (hatched gray area) up to a critical point, beyond which the capsid undergoes structural failure. Stress and strain values at the yield and critical points are indicated. The blue trace represents the window-averaged, smoothed profile derived from the raw data. (C-D) Scatter plots depicting a strong positive correlation between capsid compressive strength — defined as the stress at the critical point — and the CypA:CA bound ratio, and (D) a strong negative correlation between the critical strain — defined as the strain at the critical point— and the CypA:CA bound ratio, across all tip positions (tip position 1 to 3 in Figure 4, Supplementary Figure 5A) probed during fast AFM nanoindentation. Increased compressive strength combined with reduced tolerance to deformations indicates increased capsid brittleness as a function of the CypA:CA bound ratio. For each tip position and stoichiometry, five independent simulations were conducted (n=5); error bars represent the standard deviation.

### Cyclophilin A binding increases capsid brittleness in a stoichiometry dependent manner

To characterize the mechanical response of CypA-decorated capsids beyond the elastic regime, we performed AFM nanoindentation simulations at an increased tip velocity (v = 312.5 nm/µs; Supplementary Table 2, ID 8-14) to induce structural failure. Fast nanoindentations at the narrow end of the capsid were excluded, as rapid compression at this site caused condensation of the capsid against the AFM plate before rupture events could be captured. In this context, ductility refers to the ability of the capsid to accommodate deformation prior to failure, whereas brittleness describes mechanical failure occurring at low strain with limited deformations. Mechanical regimes, as well as yield and critical points, were identified from stress-strain profiles derived from these simulations. All capsids exhibited an initial elastic loading regime (Figure 5B, gray area) characterized by an approximately linear increase in stress with strain. This regime transitioned at a yielding point into a ductile deformation regime, where the slope of the stress-strain profile decreased and the capsid deformed more readily under applied stresses (Figure 5B, hatched area). Stress continued to increase until a critical point, corresponding to the maximum stress sustained by the capsid. Beyond this point, structural failure occurred and the AFM tip penetrated the capsid lattice, leading to a gradual reduction in stress with further indentation and marking a capsid breaking event (Figure 5A). Independent identification of these regimes was corroborated by tracking local curvature changes in the capsid during indentation (Supplementary Figure 6).

Visual inspection of the simulation trajectories revealed that the capsid failure proceeds gradually. Under increasing deformation, inter-capsomer interfaces in the capsid lattice failed sequentially, producing stepwise reductions in stress beyond the critical point. Plateaus in the stress-strain curves corresponded to transient stabilization by remaining intact interfaces prior to subsequent interface rupture events. Once all inter-capsomer interfaces under compression were disrupted, the AFM tip passed freely through the capsid lattice and the measured stress dropped to zero (Supplementary Figure 7). Thus, stress-strain profiles are information-rich as they encode both the mechanical response regimes and the progressive structural failure of the CA lattice leading to capsid breaking events.

Across all CypA:CA stoichiometries, we quantified the compressive strength and critical strain, defined as the stress and strain measured at the critical failure point, respectively. Increasing the CypA:CA bound ratio resulted in a consistent increase in the compressive strength across all indentation positions, yielding a strong positive correlation between compressive strength and the amount of CypA bound (Figure 5C). In contrast, critical strain progressively decreased with increasing CypA binding, as reflected by the strong negative correlation coefficients measured for all indentation positions (Figure 5D). Together, these trends indicate that increasing CypA binding causes the capsid to fail at lower deformations (decreased critical strain) while sustaining higher peak stresses (increased compressive strength), hallmarks of a brittle mechanical response. We therefore conclude that CypA binding increases HIV-1 capsid brittleness in a stoichiometry-dependent fashion.

### Cyclophilin A binding modulates brittleness of E45A and E45A/R132T capsid mutants

We next examined how CypA binding modulates the ductile response of two mutant capsids: the hyperstable E45A CA mutant and its revertant, E45A/R132T[45, 46]. We previously showed that the E45A mutation produces capsids with increased brittleness relative to WT, whereas the revertant mutant recovers WT-like elasticity[44]. At low amounts of bound CypA (CypA:CA bound ratios below 1:6), CypA-decorated E45A capsids consistently underwent structural failure at lower deformations than WT capsids, while addition of the R132T mutation partially recovered WT-like ductility (Figure 6). As observed for CypA-decorated WT capsids, E45A/R132T capsids became progressively more brittle with increasing amounts of bound CypA. In contrast, E45A capsids failed at low nanoindentation deformations across all CypA:CA stoichiometries, indicating a brittle mechanical response independent of the CypA bound fraction and suggesting E45A capsids are less sensitive to CypA induced brittleness. Notably, differences in ductility among CA variants diminished as the CypA:CA bound fraction increases. At CypA:CA stoichiometries exceeding 1:6, all CypA decorated capsids exhibit pronounced brittleness irrespective of CA sequence (Figure 6A), indicating that high levels of CypA binding dominate capsid mechanical behavior.

**Figure 6:**
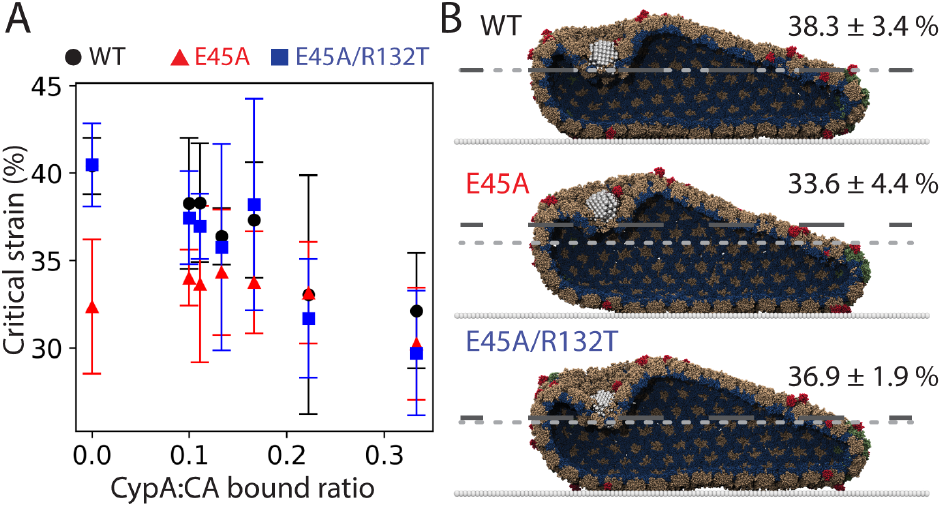
CypA-decorated capsid brittleness in the context of E45A and E45A/R132T CA mutants. (A) Scatter plot of critical strain as a function of CypA:CA bound ratio for fast AFM nanoindentation of CypA-decorated WT capsids, hyperstable E45A CA mutant capsids, and revertant E45A/R132T CA mutant capsids along the broad end (tip position 1). For each CA sequence and stoichiometry, five independent simulations were conducted (n=5); error bars represent the standard deviation. (B) Representative simulation snapshots of capsid failure events during fast AFM nanoindentation of WT, E45A and E45A/R132T CypA-decorated capsids at a 1:9 CypA:CA stoichiometry. The mean critical strain for each CA sequence is indicated by a dark grey dashed line, while the light gray dotted line denotes the WT CA critical strain baseline. SBCG2 beads are shown as van der Waals spheres, with CypA in red, CA hexamer NTD in tan, CA pentamer NTD in green, CA CTD in dark blue and the AFM tip and plate in silvery white. One half of the capsid is removed to visualize the internal structure.

## Discussion

HIV-1 nuclear entry depends on both interactions between the capsid with host cell factors as well as the intrinsic physical properties of the capsid itself. Here, we demonstrate that binding of host factor CypA modulates the HIV-1 capsid mechanoelastic response in a stoichiometry-dependent manner. Using AFM nanoindentation simulations combined with a computational framework to construct CypA-decorated capsids, we characterized capsid response across elastic, ductile and mechanical failure regimes. Although CypA binding had only modest effects on elastic stiffness, fast nanoindentation simulations revealed that CypA binding progressively increases capsid brittleness, characterized by increased compressive strength and reduced tolerance to deformations. Together, these findings establish a direct link between CypA binding and capsid brittleness, a key mechanical determinant of nuclear entry.

Our previous work demonstrated that capsid elasticity correlates with nuclear entry and infectivity: intrinsically brittle capsids, including the hyperstable E45A mutant, are defective in nuclear import, whereas capsids with WT-like elasticity, such as the E45A/R132T revertant, recover nuclear entry and infectivity[44]. The present results extend this framework by showing that CypA binding modulates capsid brittleness across WT and mutant CA sequences. WT and E45A/R132T capsids exhibited a clear brittleness increase as a function of CypA binding. E45A capsids are intrinsically brittle and, although less sensitive to CypA-induced modulation, become increasingly brittle at high CypA:CA bound fractions. These findings are consistent with recent experimental AFM observations showing that increasing CypA concentration reduces the elasticity of native HIV-1 cores and inhibits nuclear entry of intrinsically brittle capsid mutants[68].

Previous work reported that the stiffness of CA assemblies progressively increases with CypA binding, although decreasing at high CypA:CA molar ratios[47]. In contrast, we observe only a weak correlation between CypA:CA binding ratio and stiffness. Instead, stiffness correlated more strongly with local curvature of the capsid lattice. CA assemblies co-assembled with CypA exhibit increased curvature at increasing concentrations of CypA[47], therefore, apparent stiffness increases may arise from CypA-induced changes in CA assembly geometry rather than direct mechanical reinforcement of the CA lattice. Notably, our prior work showed that stiffness does not correlate with nuclear entry or infectivity[44]. Together, these observations indicate that CypA-modulated brittleness, rather than stiffness, is the mechanically relevant variable for nuclear import.

Our simulations further reveal that the HIV-1 capsid is mechanically heterogeneous, with stiffness and Poisson’s ratio varying according to local CA lattice curvature. Such mechanical heterogeneity is common in biological materials[69, 70, 71, 72] and may have functional implications for nuclear entry. Increasing evidence indicates that the HIV-1 capsid enters the cellular nucleus intact or nearly intact [14, 13], while the NPC acts as a morphology-selective filter, with preference for smaller sized capsids [73, 74], and is able to undergo deformations or even fractures [75] to accommodate capsid passage. Capsid regions of lower stiffness and high transverse deformability may facilitate progressive deformation of the capsid during NPC passage while preserving internal volume and protecting the viral genome, enhancing the likelihood of intact capsid nuclear import. Given the influence of curvature in capsid mechanoelastic properties, it is of interest to explore how different CA assembly morphologies, such as T=1 and T=4 icosahedral shaped capsids [76, 77, 78] and CA tubes [79, 80], affect the mechanoelastic response of the capsid.

Both CA mutations and CypA-binding can increase capsid brittleness and consequently impair nuclear entry. Notably, differences in ductility across CA sequences diminish at high CypA:CA binding ratios. Above a threshold of approximately 1:6 CypA:CA (one CypA molecule per CA hexamer), all capsids exhibit pronounced brittleness independent of CA sequence, indicating that CypA binding becomes the dominant determinant of capsid brittleness. This observation aligns with experimental evidence that excessive CypA binding can impair HIV-1 infection. The AC-1 CA mutant, which exhibits increased CypA-capsid binding affinity[41], shows reduced infectivity in multiple cells types, including primary CD4+ T cells[81], and can be rescued by reducing cellular expression of CypA, or by preventing CypA binding with Cyclosporin A (CsA) or CypA inhibitors[41, 37]. Cell-type specific differences in CypA expression levels further support a CypA stoichiometry-dependent effect on viral infectivity. CA mutants resitant to CypA inhibition (including A224E, A92E and G94D) replicate in Jurkat T cells, but are inhibited in H9 T cells, which express higher levels of CypA, and require CsA-mediated CypA inhibition to restore replication. In contrast, CA mutants with reduced CypA affinity are impaired in Jurkat cells but remain replication-competent in H9 cells[82, 83]. These observations suggest that CypA mediated enhancement of early replication events requires a delicate balance of the CypA fraction bound to the capsid. Together, these results support a stoichiometry-dependent mechanical model of CypA function. Partial CypA decoration modulates capsid nuclear entry by protecting the capsid from antiviral restriction factors in the cytosol while preserving sufficient capsid ductility for NPC engagement and nuclear entry. In contrast, excessive CypA binding renders capsids overly brittle and in turn impairs nuclear entry (Figure 7). Our simulations predict a critical CypA:CA binding ratio threshold about 1:6 CypA:CA, beyond which capsid brittleness becomes incompatible with nuclear entry across the CA variants presented.

**Figure 7:**
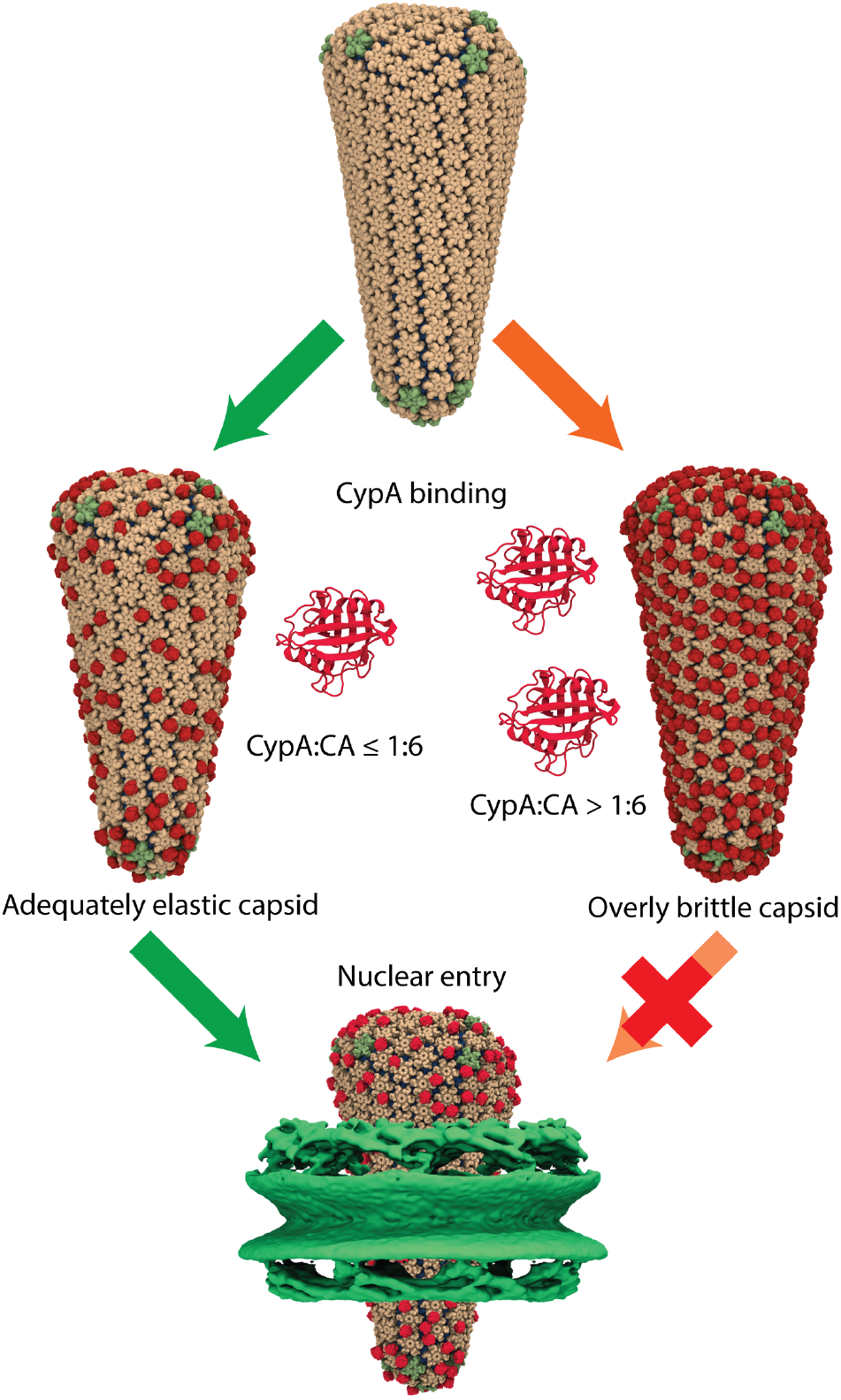
Model of CypA-mediated regulation of capsid nuclear entry. As the HIV-1 capsid transits the cytoplasm, cytoplasmic CypA binds CA at low CypA:CA ratios, stabilizing the capsid and protecting it from host restriction factors. Partial CypA decoration preserves an adequate level of capsid elasticity and permits efficient nuclear entry. In contrast, CypA decoration at high CypA:CA binding fractions, for instance, due to increased CypA-capsid binding affinity, result in overly brittle capsids that are restricted in nuclear import. CypA-decorated capsids are shown using the same color scheme as Figure 2. The human nuclear pore complex is depicted in green using a cryo-EM density map (EMD-12814).

Recent evidence suggests that CypA-mediated effects during nuclear entry are cooperative with the host factor CPSF6 within the cellular nucleus. Although CypA binds to the capsid in the cytoplasm, it potentially dissociates during nuclear entry, as suggested by differences in cryo-EM densities observed for capsids in the cytoplasm and inside the nucleus [75]. Within the nucleus, the capsid is bound by CPSF6, which modulates capsid uncoating and directs the intranuclear localization of viral integration [84, 85]. Notably, rescue of nuclear entry for capsid mutants with increased CypA binding affinity via disruption of CypA binding is dependent on CPSF6[81], indicating a functional coupling between cytoplasmic CypA binding and subsequent CPSF6 binding in the nucleus. While our current models explicitly include only CA and CypA, extending this framework to incorporate CPSF6 and the viral genome will be essential for understanding how coordinated capsid-host factor interactions regulate nuclear entry, uncoating dynamics and vDNA delivery inside the nucleus.

Overall, our findings reinforce the emerging view that capsid mechanoelastic properties are critical determinants of HIV-1 nuclear entry and infectivity [44, 73]. By establishing a direct mechanistic link between CypA binding and capsid brittleness, this work provides a physical framework for understanding how host cell factors regulate viral replication through modulation of the capsid mechanoelastic response.

## Acknowledgements

The authors acknowledge funding from the US National Institutes of Health awards R01AI178846 and U54AI170791 (to J.R.P). This work used the Stampede 3 supercomputer at the Texas Advanced Supercomputing Center and the Delta advanced computing system at the National Center for Supercomputing Applications and the University of Illinois Urbana-Champaign through allocation MCB170096 from the Advanced Cyberinfrastructure Coordination Ecosystem: Services & Support (ACCESS) program, which is supported by U.S. National Science Foundation grants #2138259, #2138286, #2138307, #2137603, and #2138296.

## Data Availability

All scripts, input structures and parameters necessary for conducting the simulations described in this manuscript, as well as example outputs and the necessary scripts for analyzing the simulation trajectories are deposited in a publicly available Zenodo repository (https://doi.org/10.5281/zenodo.17653432).

## Author contributions

This research was conceived by J.R.P. J.R.P. acquired funding for this research. A.J.B. constructed and parameterized shape-based coarse-graining models of CypA. Modeling of CypA-decorated capsids was performed by J.S.R. J.S.R. and A.J.B. conducted MD simulations. J.S.R. performed analyses and drafted figures under J.R.P. guidance. J.S.R. and J.R.P wrote the original draft of the manuscript.

