## Supplementary Information for "Stoichiometric binding of Cyclophilin-A to the HIV-1 capsid modulates its mechanoelastic properties"

Juan S. Rey[1] Alexander J. Bryer[1] Juan R. Perilla[1,*]

February 20, 2026

[1]Department of Chemistry & Biochemistry, University of Delaware, Newark, DE 19716

**This PDF includes:**
Supplementary Table 1 and 2
Supplementary Figure 1 to 7

Supplementary Table 1: CypA-decorated capsid CypA:CA stoichiometries and CypA:CA binding ratios modeled in this study. Each CypA-decorated capsid model consists of 241 CA hexamers and 12 pentamers (1506 CA monomers) and a variable number of CypA molecules. A CypA:CA stoichiometry of "None" denotes an undecorated capsid.

| **CypA:CA** | **CypA:CA binding ratio** | **No. CA** | **No. CypA** |
|---|---|---|---|
| None | 0.0 | 1506 | 0 |
| 1:10 | 0.10 | 1506 | 151 |
| 1:9 | 0.11 | 1506 | 167 |
| 1:7.5 | 0.13 | 1506 | 201 |
| 1:6 | 0.17 | 1506 | 251 |
| 1:4.5 | 0.22 | 1506 | 335 |
| 1:3 | 0.33 | 1506 | 502 |

Supplementary Table 2: Summary of AFM nanoindentation simulations. Slow nanoindentation simulations (ID 1-7) were performed with AFM tip velocity of 3.125 nm/µs for WT CypA-decorated capsids at tip positions 1 to 4. Fast nanoindentation simulations (ID 8-28, tip velocity 312.5 nm/µs) were performed for WT, E45A, and E45A/R132T capsids at tip positions 1 to 3.

| Simulation ID | CA sequence | CypA:CA stoichiometry | Probe vel. (nm/µs) | Simulation length (ns) | Tip pos. probed | Num. replicas |
|---|---|---|---|---|---|---|
| 1 | WT | None | 3.125 | 20,000 | 1-4 | 3 |
| 2 | | 1:10 | 3.125 | 20,000 | 1-4 | 3 |
| 3 | | 1:9 | 3.125 | 20,000 | 1-4 | 3 |
| 4 | | 1:7.5 | 3.125 | 20,000 | 1-4 | 3 |
| 5 | | 1:6 | 3.125 | 20,000 | 1-4 | 3 |
| 6 | | 1:4.5 | 3.125 | 20,000 | 1-4 | 3 |
| 7 | | 1:3 | 3.125 | 20,000 | 1-4 | 3 |
| 8 | WT | None | 312.500 | 250 | 1-3 | 5 |
| 9 | | 1:10 | 312.500 | 250 | 1-3 | 5 |
| 10 | | 1:9 | 312.500 | 250 | 1-3 | 5 |
| 11 | | 1:7.5 | 312.500 | 250 | 1-3 | 5 |
| 12 | | 1:6 | 312.500 | 250 | 1-3 | 5 |
| 13 | | 1:4.5 | 312.500 | 250 | 1-3 | 5 |
| 14 | | 1:3 | 312.500 | 250 | 1-3 | 5 |
| 15 | E45A | None | 312.500 | 250 | 1-3 | 5 |
| 16 | | 1:10 | 312.500 | 250 | 1-3 | 5 |
| 17 | | 1:9 | 312.500 | 250 | 1-3 | 5 |
| 18 | | 1:7.5 | 312.500 | 250 | 1-3 | 5 |
| 19 | | 1:6 | 312.500 | 250 | 1-3 | 5 |
| 20 | | 1:4.5 | 312.500 | 250 | 1-3 | 5 |
| 21 | | 1:3 | 312.500 | 250 | 1-3 | 5 |
| 22 | E45A/R132T | None | 312.500 | 250 | 1-3 | 5 |
| 23 | | 1:10 | 312.500 | 250 | 1-3 | 5 |
| 24 | | 1:9 | 312.500 | 250 | 1-3 | 5 |
| 25 | | 1:7.5 | 312.500 | 250 | 1-3 | 5 |
| 26 | | 1:6 | 312.500 | 250 | 1-3 | 5 |
| 27 | | 1:4.5 | 312.500 | 250 | 1-3 | 5 |
| 28 | | 1:3 | 312.500 | 250 | 1-3 | 5 |

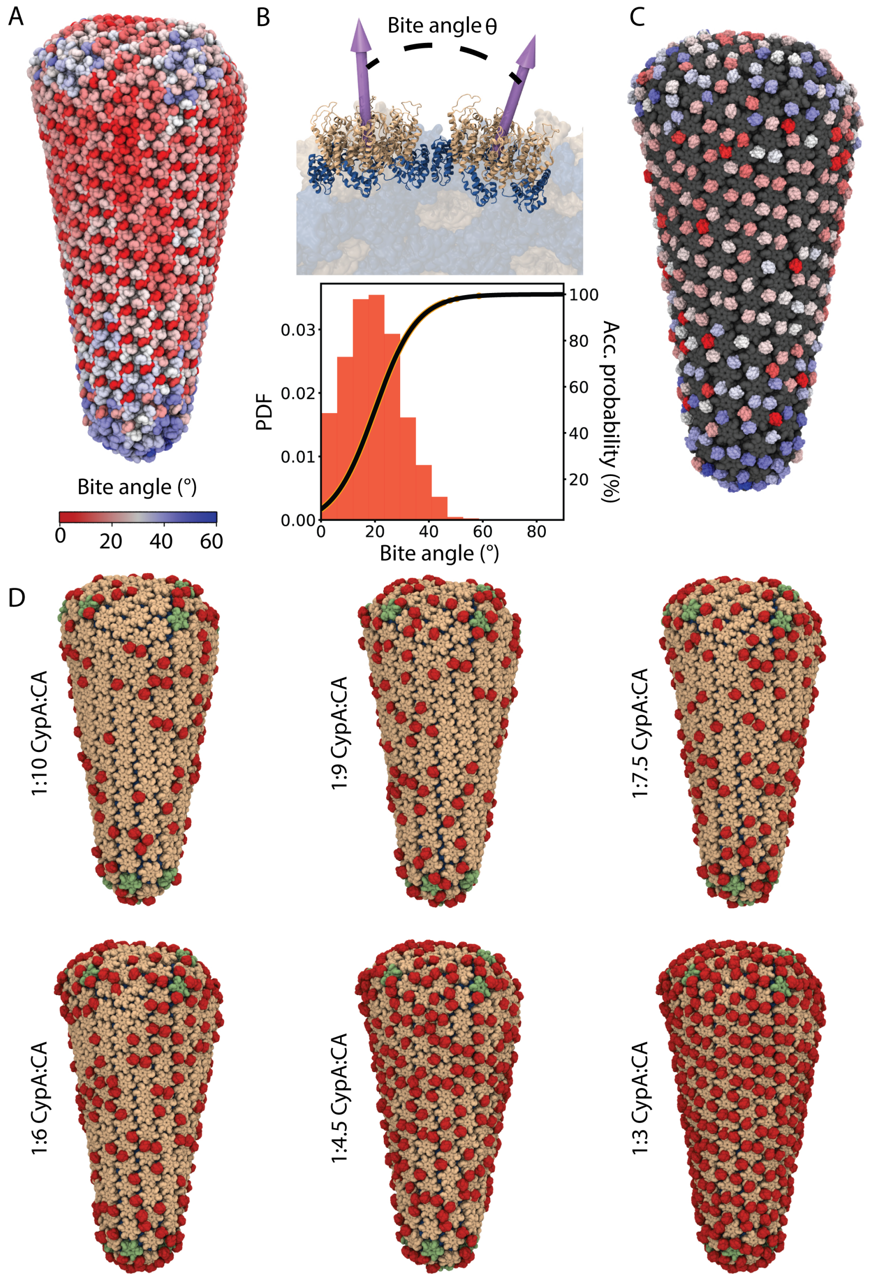


Supplementary Figure 1: Curvature-based construction of CypA-decorated capsid models. (A) All-atom HIV-1 capsid model. Each CA dimer is colored according to the bite angle between the capsomers it connects. (B) (Top) Schematic illustrating bite angle calculation between two adjacent capsomers. The bite angle is defined as the change in orientation between the director vectors of neighboring capsomers. Two CA hexamers are shown in cartoon representation, with CA NTD and CTD colored tan and blue, respectively; capsomer director vectors are shown as purple arrows. (Bottom) Bite angle distribution for all adjacent capsomer pairs in the all-atom capsid model (orange bars), overlaid with the curvature-biased probability density function used in the random-walk CypA decoration procedure (solid black line). (C) Fully saturated CypA-decorated capsid model generated using the random-walk algorithm. The capsid molecular surface is colored in gray, and CypA molecules are colored according to the bite angle of the CA dimers at their binding interfaces. (D) CypA-decorated capsid models used for the simulations in this study, spanning CypA:CA stoichiometries from 1:3 to 1:10 (see Table 1). CypA molecules are colored in red, and the capsid molecular surface is colored dark blue (CA-CTD), tan (CA hexamer NTD) and green (CA pentamer NTD).

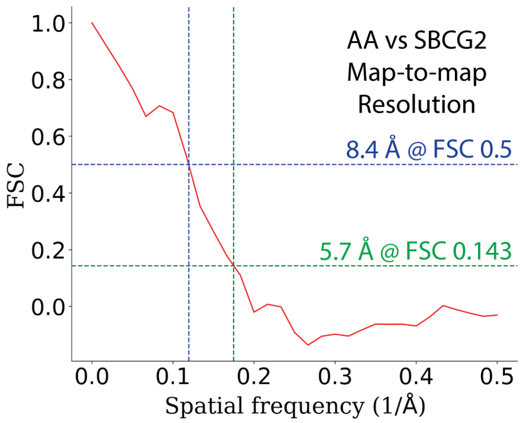


Supplementary Figure 2: Representative FSC curve comparing AA and SBCG2 charge distributions for a CypA SBCG2 model with 165 beads. The effective charge resolution is estimated to be 8.4 Å and 5.7 Å at the FSC thresholds 0.5 and 0.143, respectively.

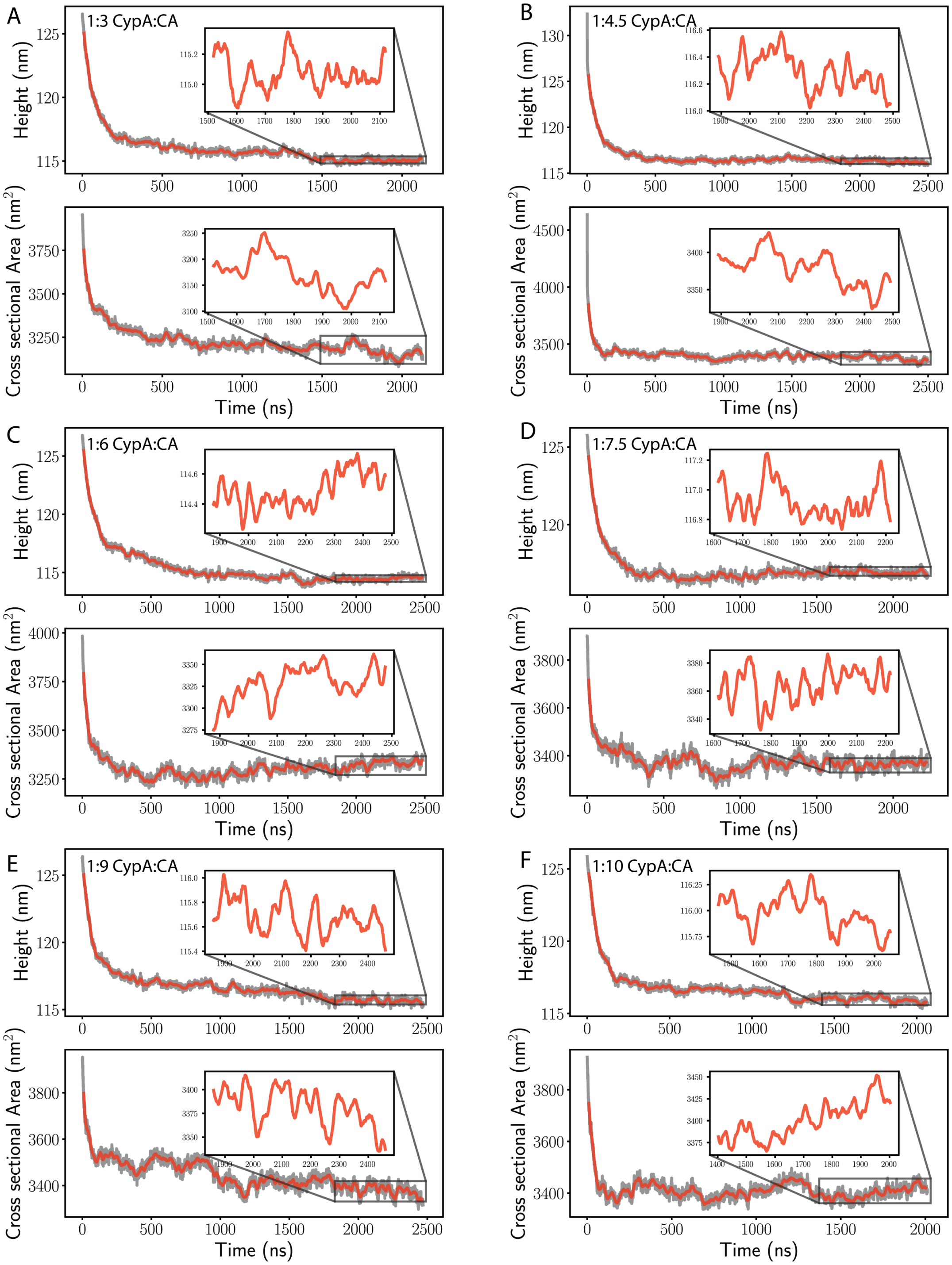


Supplementary Figure 3: Equilibration of CypA-decorated HIV-1 capsids. Capsid height and cross-sectional area during equilibration of CypA-decorated capsids at CypA:CA stoichiometries of (A) 1:3, (B) 1:4.5, (C) 1:6, (D) 1:7.5, (E) 1:9, and (F) 1:10. Raw data points are shown in gray, and windowed-averaged traces are shown in red. Insets show convergence of capsid height and cross-sectional area fluctuations to below 1% over 2 μs equilibration trajectories.

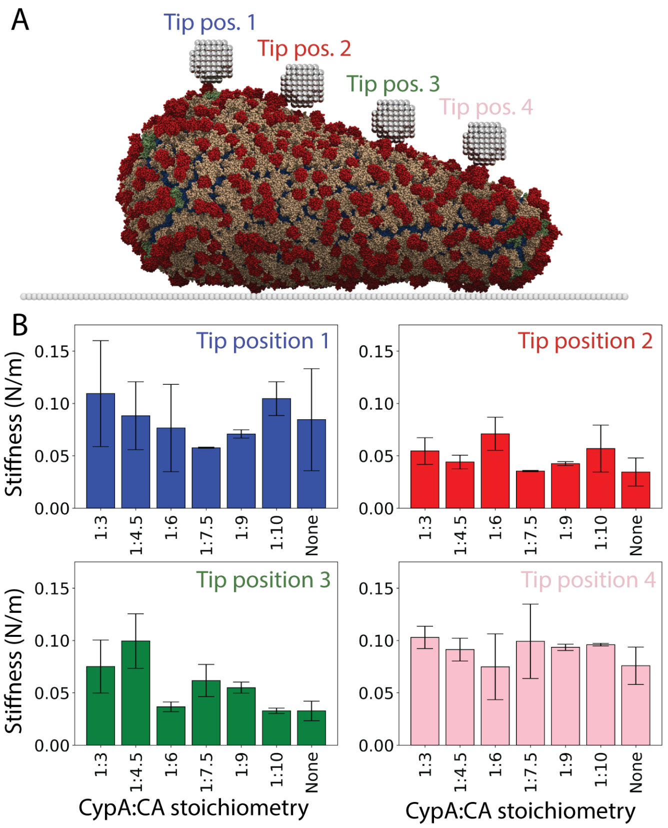


Supplementary Figure 4: Capsid stiffness measured at distinct locations along the capsid surface. (A) Side view of a CypA-decorated capsid at 1:3 CypA:CA stoichiometry in the AFM setup. The capsid is colored in tan (CA NTD), blue (CA CTD), green (CA pentamer NTD) and red (CypA), while the AFM plate and tip are depicted using gray and silver beads. The four tip positions utilized in the slow AFM nanoindentation simulations are indicated with the AFM tip and labeled. (B) Bar plots for the mean stiffness measured during AFM nanoindentations along all tip positions for CypA-decorated capsids across our CypA:CA stoichiometry range. For each stoichiometry and tip position three independent simulation replicas were conducted (n=3); error bars represent standard deviation.

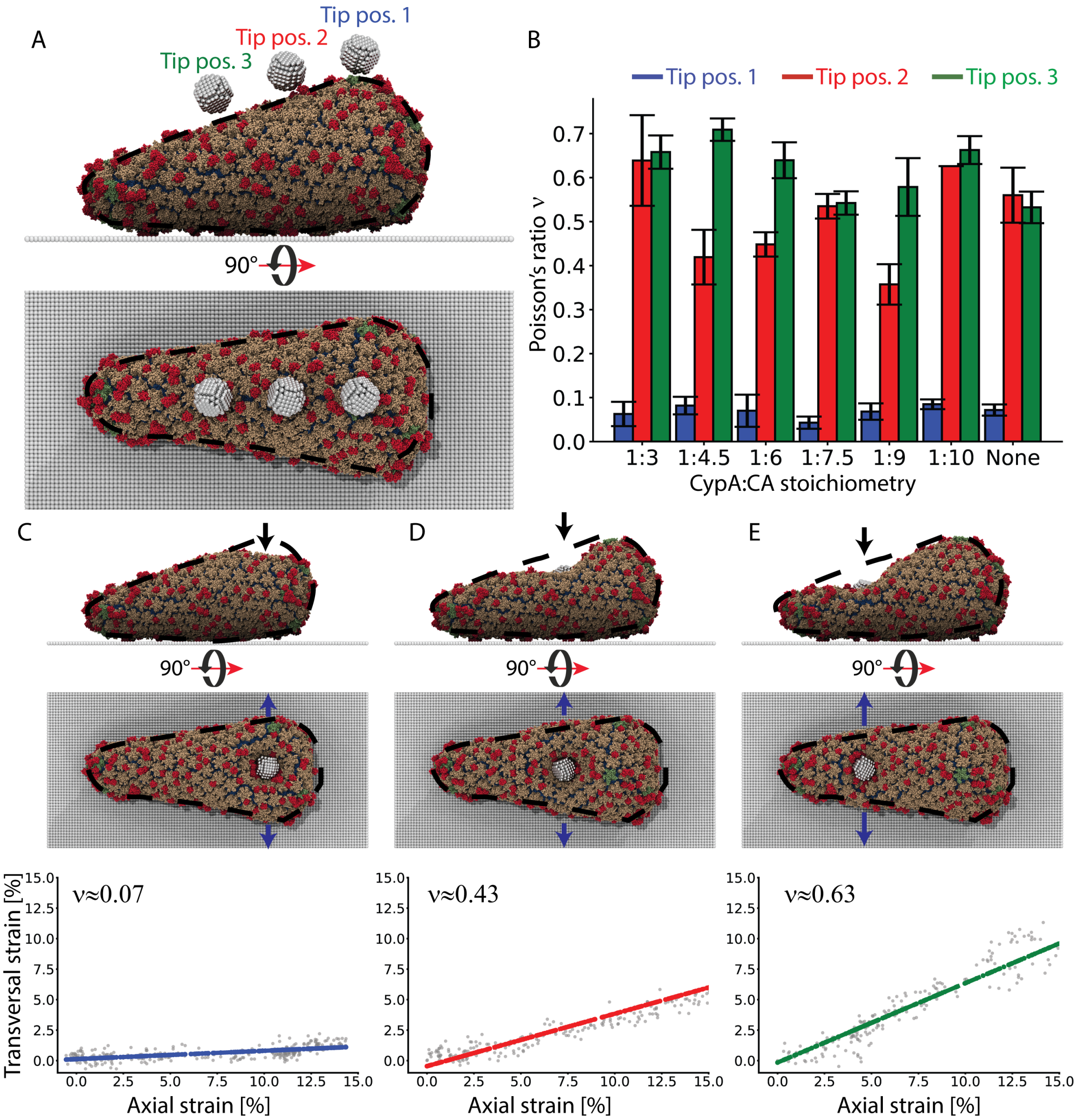


Supplementary Figure 5: Poisson's ratio characterization at different regions of the HIV-1 capsid. (A) Rendering of a CypA-decorated capsid at 1:6 CypA:CA stoichiometry in the fast AFM nanoindentation setup. In each simulation, the AFM tip is positioned above one of three locations: the broad end of the capsid (tip position 1) or two flatter regions along the upper (tip position 2) or lower (tip position 3) halves of the capsid. The SBCG CypA-decorated capsid is shown using the color scheme of Figure 5A; AFM tip and plate are shown in metallic and opaque gray, respectively. (B) Bar plot showing the mean Poisson's ratio calculated from fast AFM nanoindentation simulations at the tip positions shown in A, across the established range of CypA:CA stoichiometries (see Supplementary Table 1). For each tip position and stoichiometry, five independent nanoindentation simulations were performed (n=5); error bars represent standard deviation. (C-E) (Top) Top and side view snapshots illustrating axial and transversal deformation of a CypA-decorated capsid at 1:6 CypA:CA stoichiometry during AFM nanoindentation at tip positions 1 (C), 2 (D), and 3 (E). Dashed black lines denote the silhouette of the capsid before compression, black and blue arrows indicate the directions of axial strain (compression) and transversal strain (expansion), respectively. (Bottom) Representative axial vs. transversal strain traces. Raw data points are shown in gray, and linear fits are shown in blue, red or green, corresponding to the tip position labels in B. Poisson's ratios $\nu$ are calculated from the slopes of linear fits.

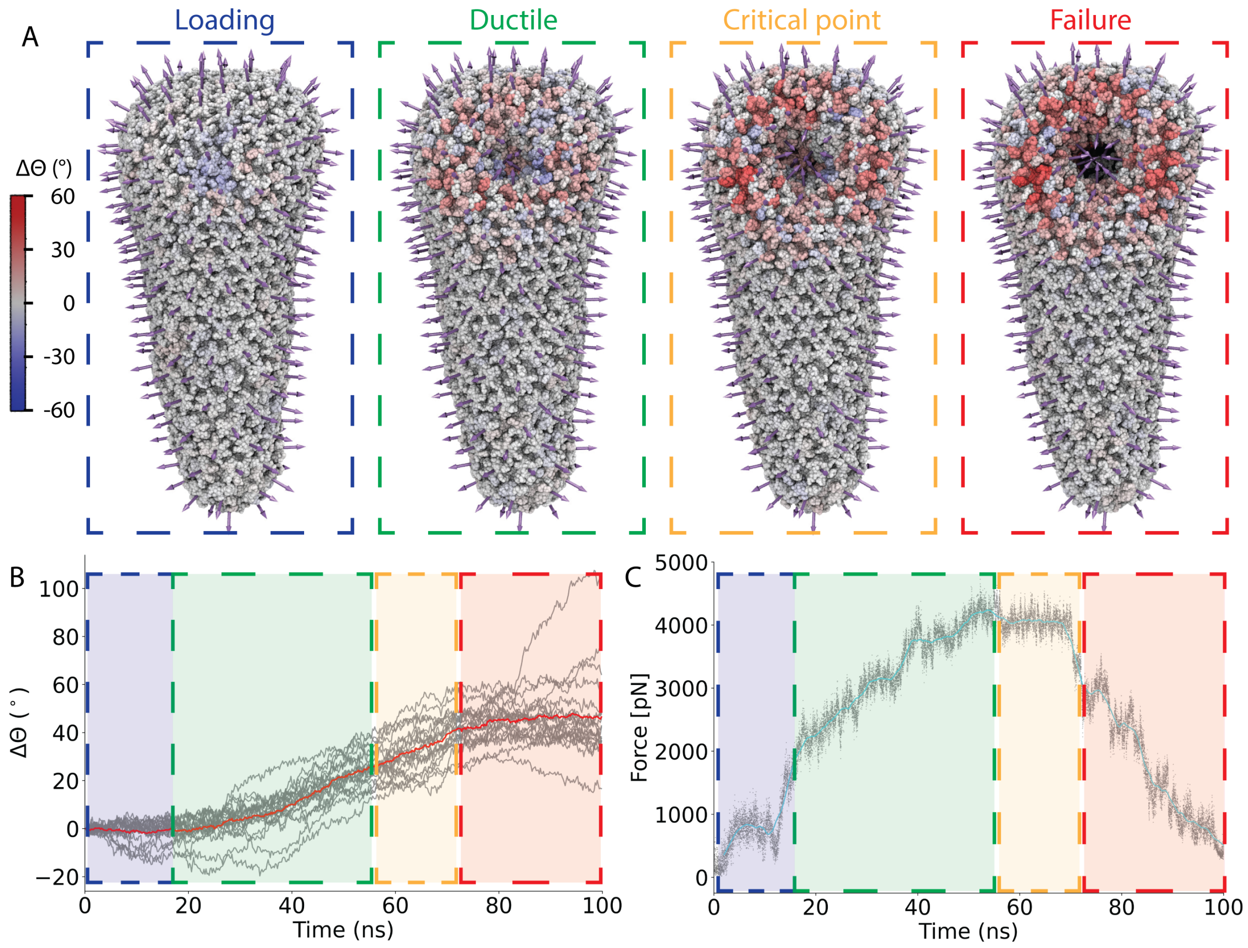


Supplementary Figure 6: Curvature-based characterization of capsid mechanical regimes. (A) Simulation snapshots of a CypA-decorated capsid at 1:3 CypA:CA stoichiometry (CypA molecules omitted for visualization clarity) at different mechanical regimes during nanoindentation. CA dimers are colored according to the change in the bite angle ($\Delta\Theta$) between the capsomers they connect, relative to the undeformed capsid structure before nanoindentation. Capsomer director vectors are shown as purple arrows. Snapshots correspond to the loading regime (blue box), characterized by minimal curvature changes; the ductile deformation regime (green box), marked by localized curvature changes; the critical point (yellow box), where applied stress is maximal; and during the structural failure regime (red box). (B) Bite angle change traces for capsomers neighboring the nanoindentation axis (gray) and the average bite angle change (red). Dashed boxes indicate the distinct mechanical regimes using the same color scheme as panel A. (C) Corresponding force-time nanoindentation profile. Raw data points are colored in gray, and the windowed average trace is shown in cyan. Mechanical regimes are indicated using dashed colored boxes, consistent with panel A.

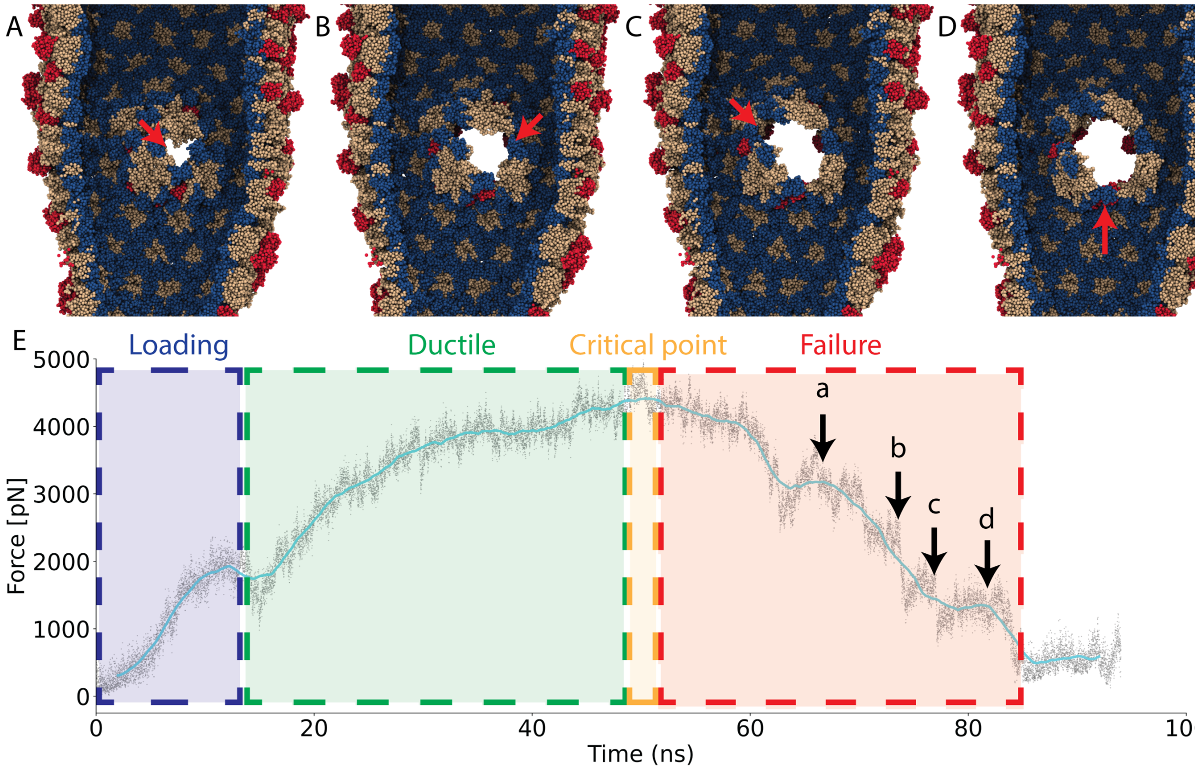


Supplementary Figure 7: Capsid structural failure is a gradual process involving sequential disruption of CA interfaces. (A-D) Sequential simulation snapshots illustrating the internal structural failure of a CypA-decorated capsid at 1:3 CypA:CA stoichiometry during AFM nanoindentation. (A) Initial disruption of the CA dimer interfaces the two hexamers closest to the AFM tip is followed by sequential separation of neighboring capsomers and disruption of additional dimer interfaces (B-D), ultimately resulting in complete structural failure. The SBCG2 CypA-decorated capsid is colored according to the color scheme in Figure 5A. Disrupted dimer interfaces are indicated by red arrows. (E) Corresponding force-time AFM nanoindentation trace, with mechanoelastic regimes labeled as in Supplementary Figure 6. Distinct force plateaus observed in the failure regime correspond to the resistance associated with disruption of individual CA dimer interfaces and are indicated with black arrows labeled a-d, corresponding to the snapshots in panels A-D.
